# Designing safe and potent herbicides with the cropCSM online resource

**DOI:** 10.1101/2020.11.01.364240

**Authors:** Douglas E. V. Pires, Keith A. Stubbs, Joshua S. Mylne, David B. Ascher

## Abstract

Herbicides have revolutionised weed management, increased crop yields and improved profitability allowing for an increase in worldwide food security. Their widespread use, however, has also led to not only a rise in resistance but also concerns about their environmental impact. To help identify new, potent, non-toxic and environmentally safe herbicides we have employed interpretable predictive models to develop the online tool cropCSM (http://biosig.unimelb.edu.au/crop_csm).

---

Developing herbicides, much like pharmaceuticals, involves a careful balance between efficacy and safety. In the pharmaceutical industry, drug development pipelines have tackled these challenges by modelling and optimising these important parameters early in the development process. This has led, in general, to increased hit rates and decreased attrition due to poor toxicity profiles and, in the process, reduced development time, costs, and animal testing^1–4^. Although many computer-guided approaches have proven invaluable for drug development, by contrast little has been done to aid the development of safe and potent agrochemicals.

Using experimental information on the herbicidal activity of over 4,000 small molecule compounds (22% with herbicidal activity), we investigated what physicochemical properties of the compounds translate to herbicidal activity. Herbicidal molecules were enriched in saturated carbon chains and benzene substructures, compared to the inactive molecules **(Fig. 1a).** The majority (90%) of the active compounds tended to be less than 517 Da, up to 9 acceptors and 4 donors, with fewer than 9 rotatable bonds and a logP between −1.7 and 6.1 **(Supplementary Fig. 1**) (95% less than 700 Da, 11 rotatable bonds, 11 acceptors, 6 donors, and logP −3.0 to 6.1). This is similar, although slightly more lenient, than the widely used Lipinski Rule of Five for orally bioavailable drugs. Interestingly, but consistently, there was no significant distinction in physicochemical properties between herbicides and approved drugs, as illustrated in the t-SNE plot **(Supplementary Fig. 2).** Compared to all FDA approved drugs, however, herbicides were enriched in substructures involving chlorine.

**Figure 1.**
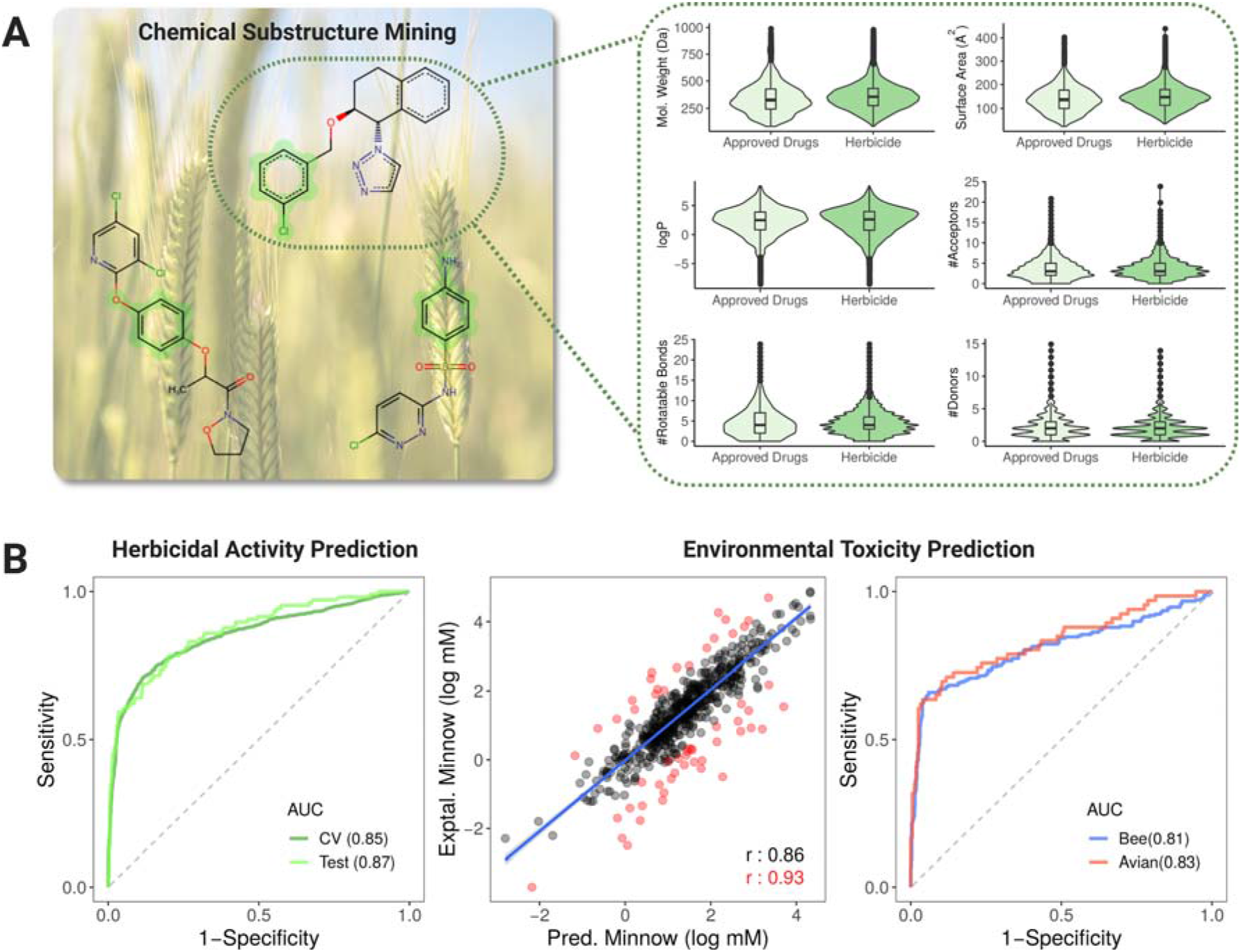
cropCSM: predicting safe and potent herbicides. Using chemical substructure mining we identified common enriched substructures in compounds with herbicidal activity (A-left). Active compounds presented similar molecular properties of approved drugs (A-right). Performance of herbicide and environmental-toxicity predictors is shown in (B). Our herbicide predictor was able to accurately identify active compounds with AUC>0.85 on cross-validation and blind test. Three environmental toxicity models have been developed and were capable of successfully measuring minnow toxicity (as a regression task, center graph) as well as identifying potentially harmful compounds for Bees and Mallard (right-hand side graph).

These insights were used as a platform to build a supervised machine learning predictive model, where the small molecule structure was represented as a graph-based signature, termed Cutoff Scanning Matrix (CSM, in which the atoms are represented as nodes, and covalent interactions between them as edges^5,6^ **(Supplementary Fig. 3).** Under crossvalidation, we were able to correctly identify 82% of the active molecules with an overall accuracy of 87% and AUC of 0.85 **(Fig. 1b** and **Supplementary Table 1**). When the model was evaluated against a blind test set of 106 active and 345 inactive molecules, we achieved comparable performance (87% accuracy, AUC of 0.87). This provided confidence that the approach can be generalized and used with unknown sets of putative herbicidal molecules active against a target of interest.

Agrochemicals have been linked to a range of unwanted negative effects on both health and the environment. To help identify safe herbicides, complementary models were developed to capture the impact of a small molecule on the honey bee *(Apis mellifera),* mallard *(Anas platyrhynchos)* and flathead minnow *(Pimephales promelas)* toxicity, in addition to measures of human health, including AMES toxicity, rat LD_50_ and oral chronic toxicity. While assessing molecular substructures enriched in toxic compounds, **(Supplementary Fig. 4),** we identified a prevalence of complex ring structures. Of note, structures rich in chlorine, while enriched in herbicides, were also enriched in compounds that were toxic for mallard and minnow, highlighting a potential inherent difficulty in optimising potency and safety when designing herbicides.

We were also able to identify toxic molecules as classification and regression tasks with accuracies of up to 92% and Pearson’s correlations of up to 0.86, outperforming previous predictive approaches **(Fig. 1b, Supplementary Fig. 5** and **Supplementary Tables 2-3).** These results add credence to the tool to rapidly identify potentially hazardous molecules early in the development process, which has the potential to significantly reduce costs and failure rates.

The cropCSM models were then applied to a set of 360 commercial herbicides^7^. Over 97% were correctly identified as herbicidal **(Fig. 2).** Despite being outliers in terms of their physicochemical properties, cropCSM correctly predicted glyphosate and paraquat as herbicides. Of those that weren’t, however, they included the natural fatty acid oleic acid, and non-specific fragments like molecules such as dazomet and pentachlorophenol.

**Figure 2.**
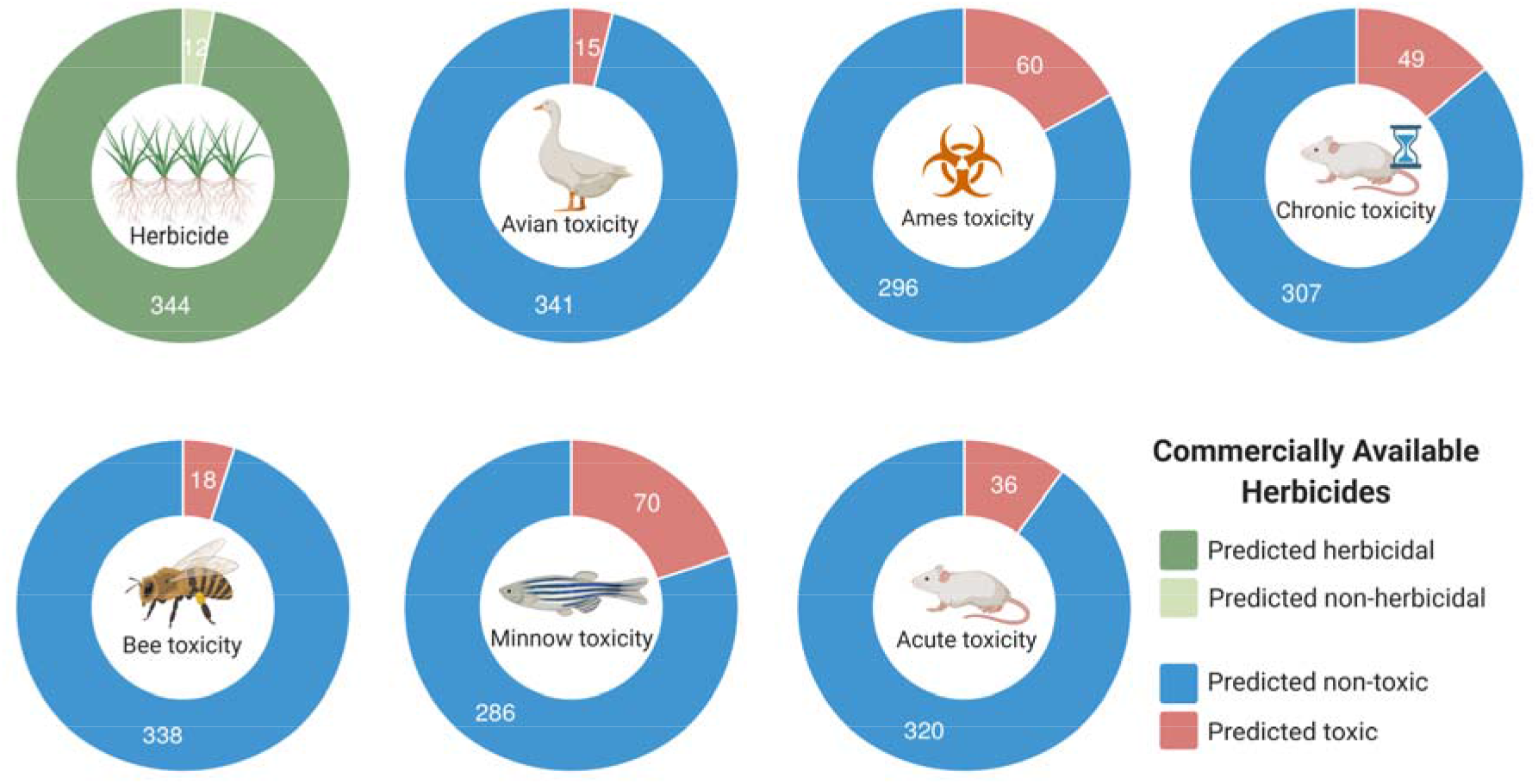
Performance of cropCSM on commercially available herbicides. Our method was able to correctly classify 97% of commercial herbicides (344 out of 356, top-left graph). The figure also shows the proportion of compounds predicted to be environmental or human toxic. Molecules were more frequently predicted as AMES toxic (17%, 60 out of 356) and Minnow toxic (20%, 70 out of 356).

Overall, our cropCSM tool provides the first free and easy-to-use *in silico* platform to help develop herbicides that are safe, effective and minimise impact on the environment. We anticipate future iterations of cropCSM that will draw upon larger datasets and as a result will have a higher predictor capability, allowing for a greater increase in accuracy and correlation. The herbicidal and toxicity predictors are freely available via an integrated and easy-to-use web interface **(Supplementary Fig. 6;** http://biosig.unimelb.edu.au/crop_csm).

## Acknowledgements

D.B.A. and D.E.V.P. were funded by the Jack Brockhoff Foundation (JBF 4186, 2016) and an Investigator Grant from the National Health and Medical Research Council (NHMRC) of Australia (GNT1174405). Supported in part by the Victorian Government’s Operational Infrastructure Support Program. The authors thank Julie Leroux, Joel Haywood, Kirill Sukhoverkov and Kalia Bernath-Levin for acquiring the 4,513 data points used to train cropCSM. To acquire the dataset, we also thank for funding the UWA Office of Industry and Innovation Pathfinder Funding Scheme, the Australian Research Council (ARC) for a Future Fellowship (FT120100013) to J.S.M., Nexgen Plants Pty Ltd and ARC grant DP190101048 to J.S.M. and K.A.S.

## Author Contributions

D.E.V.P. and D.B.A. were responsible for method design and website development. K.A.S. and J.S.M. were responsible for development of datasets on herbicidal activity. All authors assisted with manuscript writing.

## Additional Information

Supplementary Information is available.

## ONLINE METHODS

### Data for Herbicidal Activity

A dataset of 4,513 experimentally characterized, structurally diverse small molecules and their herbicidal activity profiles^7,8^. These were labelled either as active (997 molecules) or inactive (3,516 molecules). They had an average molecular weight of 380 Da and logP of 2.4 **(Supplementary Fig. 1**). A database of 360 commercial herbicides was also used to evaluate cropCSM^7,8^.

### Environmental and Human Toxicity

We have developed new predictors based on six environmental and human toxicity data sets with experimentally characterised molecules. Environmental toxicity data sets included (i) honey bee (A. *mellifera)* toxicity, which was composed of 247 toxic and 353 atoxic molecules^9^; (ii) avian toxicity, composed of 461 small molecules and their effects on mallard duck (66 toxic and 395 atoxic)^10^ and (iii) flathead minnow toxicity, with lethal concentration values (LC50) for a diverse set of 554 molecules^11^. Human toxicity data sets included (i) AMES toxicity, with compounds labelled based on their carcinogenic potential (4,632 carcinogenic and 3,470 not-carcinogenic)^12^; (ii) oral acute toxicity in rats, denoted as lethal dose (LD50) values for 10,145 compounds^13^ and (iii) oral chronic toxicity in rats values for 567 compounds^14^.

### Graph-based Signatures and Feature Engineering

Graph modelling has an invaluable tool to model biological entities, including small molecules. Over the years we have proposed and developed the concept of graph-based signatures (based on Cutoff Scanning Matrix concept^15^) to represent physicochemical and geometrical properties of a range of macromolecules^5,16–18^ and their interactions^19–25^. These have also been successfully adapted to represent small molecules pharmacokinetics, toxicity and bioactivity^5,6^. **Supplementary Fig. 3** depicts the main steps involved in feature engineering with graph-based signatures. Small molecules are modelled as unweighted, undirected graphs where nodes represent atoms and edges represent covalent bonds. Via pharmacophore modelling^5^, atoms are labelled based on their properties and allpairs shortest paths distances are calculated. Molecules are then represented as cumulative distribution functions of atom distances labelled based on their respective physicochemical properties (pharmacophores) and converted as a feature vector used as evidence to train and test predictive methods. Complementary physicochemical properties are calculated and included using the RDKit cheminformatics library^26^ and included in the feature vector. Frequent substructure mining was performed using MoSS^27^.

### Model Selection and Validation

Different supervised learning algorithms available on the scikit-learn Python library^28^ were assessed with best performing models selected based on Matthew’s Correlation Coefficient (MCC) and the Area under the ROC curve (AUC) for classification tasks and Pearson’s correlation and Root Mean Squared Error (RMSE) for regression tasks. Performance was assessed under 10-fold cross validation as well as using non-redundant blind tests. A feature selection step was used to reduce dimensionality and improve performance via a Forward Greedy Selection approach.

### Web server

The backend of the cropCSM web server was developed using the Python Flask framework version 0.12.3 and the front end using Bootstrap framework version 3.3.7. The system is hosted by a Linux server running Apache.

