## Supplementary Information for "Designing safe and potent herbicides with the cropCSM online resource"

**Supplementary Table 1.** Performance of cropCSM on identifying distinguishing molecules with herbicidal activity.

|  | **Performance metrics** | | |
| --- | --- | --- | --- |
| ***Data set*** | **Accuracy** | **AUC** | **MCC** |
| Cross-validation | 87% | 0.85 | 0.60 |
| Blind test | 87% | 0.87 | 0.59 |

**Supplementary Table 2.** Performance of cropCSM on identifying environmentally toxic compounds as a regression task. The table presents performance assessed on cross-validation.

| **Environmental Toxicity** | **Cross Validation** | |
| --- | --- | --- |
|  | **Pearson (r)** | **RMSE** |
| ***Minnow Toxicity (LC50)*** | | |
| cropCSM | 0.86 | 0.74 |
| pkCSM | 0.74* | 0.84 |
| admetSAR | 0.57* | 0.67 |
| ***Oral Rat Chronic Toxicity*** | | |
| cropCSM | 0.75 | 0.56 |
| pkCSM | 0.68* | 0.74 |
| Mazzatorta *et al.* | 0.50* | 0.73 |
| ***Oral Rat Acute Toxicity (LD50)*** | | |
| cropCSM | 0.79 | 0.62 |
| pkCSM | 0.66* | 0.68 |
| admetSAR | 0.61* | 0.32 |

*Denotes a statistically significant performance difference obtained via a Fisher r−to−z transformation, by calculating the z value, using a threshold of p ≤ 0.05 for significance.

**Supplementary Table 3.** Performance of cropCSM on identifying environmentally toxic compounds as a classification task. The table presents performance assessed on cross-validation.

| **Environmental Toxicity** | **Cross Validation** | | |
| --- | --- | --- | --- |
|  | **Accuracy** | **AUC** | **MCC** |
| ***Honey Bee Toxicity*** | | | |
| cropCSM | 0.87 | 0.81 | 0.65 |
| Wang *et al.* | 0.84 | 0.84 | 0.60 |
| ***Avian Toxicity*** | | | |
| cropCSM | 0.92 | 0.83 | 0.65 |
| Zhang *et al.* | 0.91 | 0.81 | - - - |
| ***AMES Toxicity*** | | | |
| cropCSM | 0.87 | 0.94 | 0.74 |
| pkCSM | 0.84 | 0.91 | - - - |
| admetSAR | 0.85 | 0.91 | - - - |

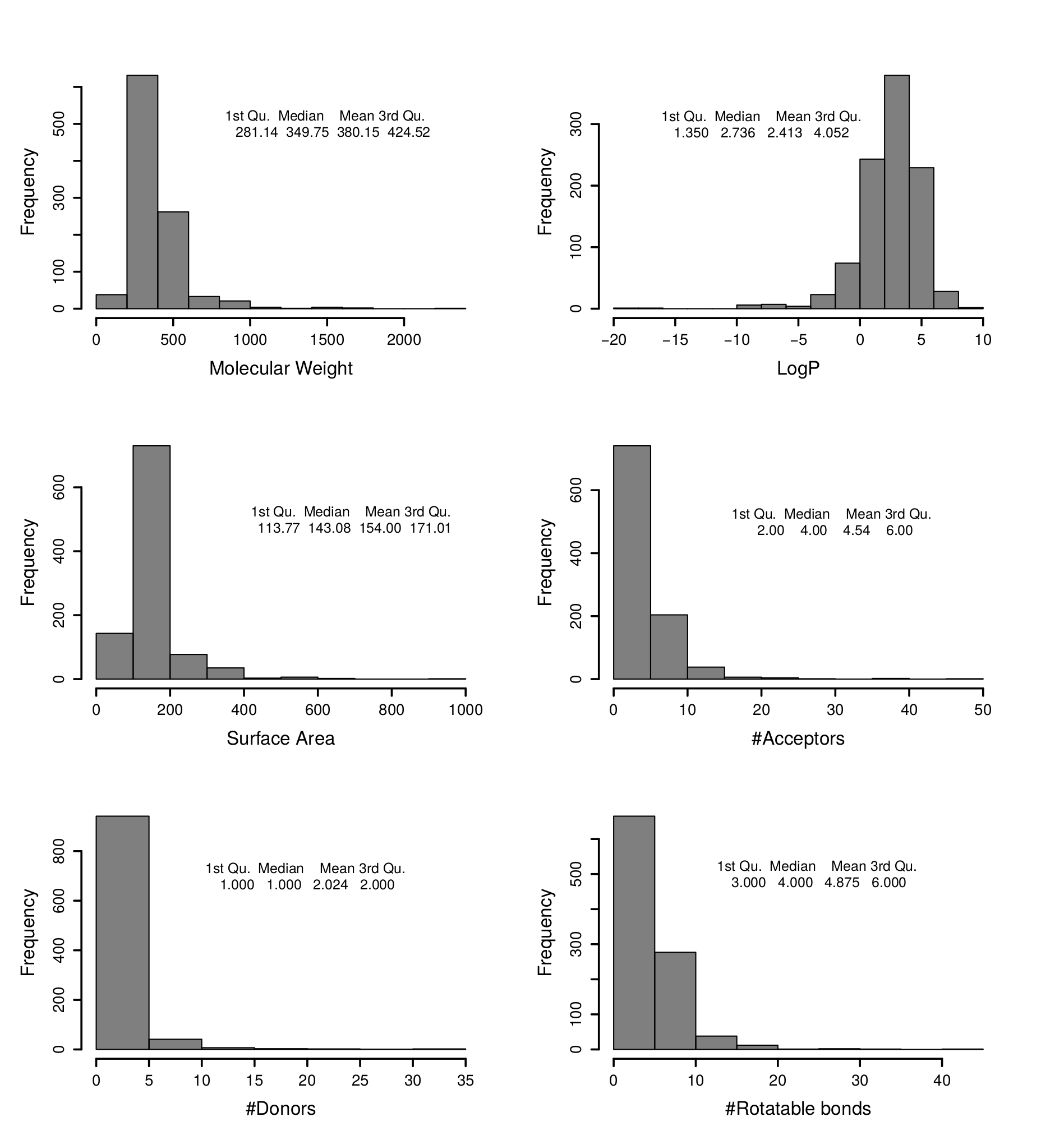

**Supplementary Fig. 1.** Property distribution for molecules with herbicidal activity. Herbicides seem to present similar properties, although slightly more lenient, to orally bioavailable drugs adherent to the Lipinski’s Rule of 5.

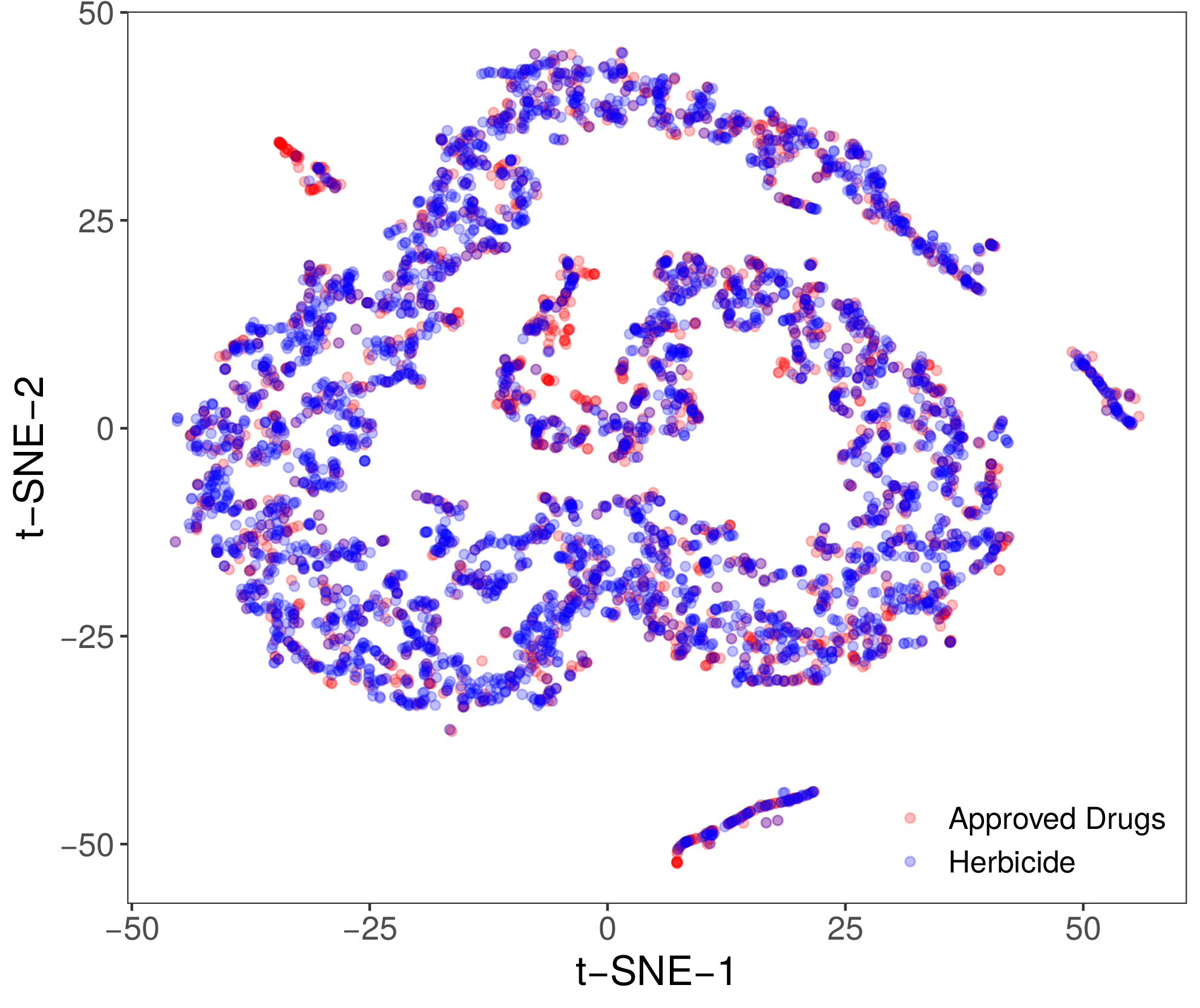

**Supplementary Fig. 2.** t-SNE plot comparing common physicochemical properties of approved drugs and herbicides. No distinction was identified between these two groups of molecules for a range of different parameters (t-SNE perplexity was systematically assessed).

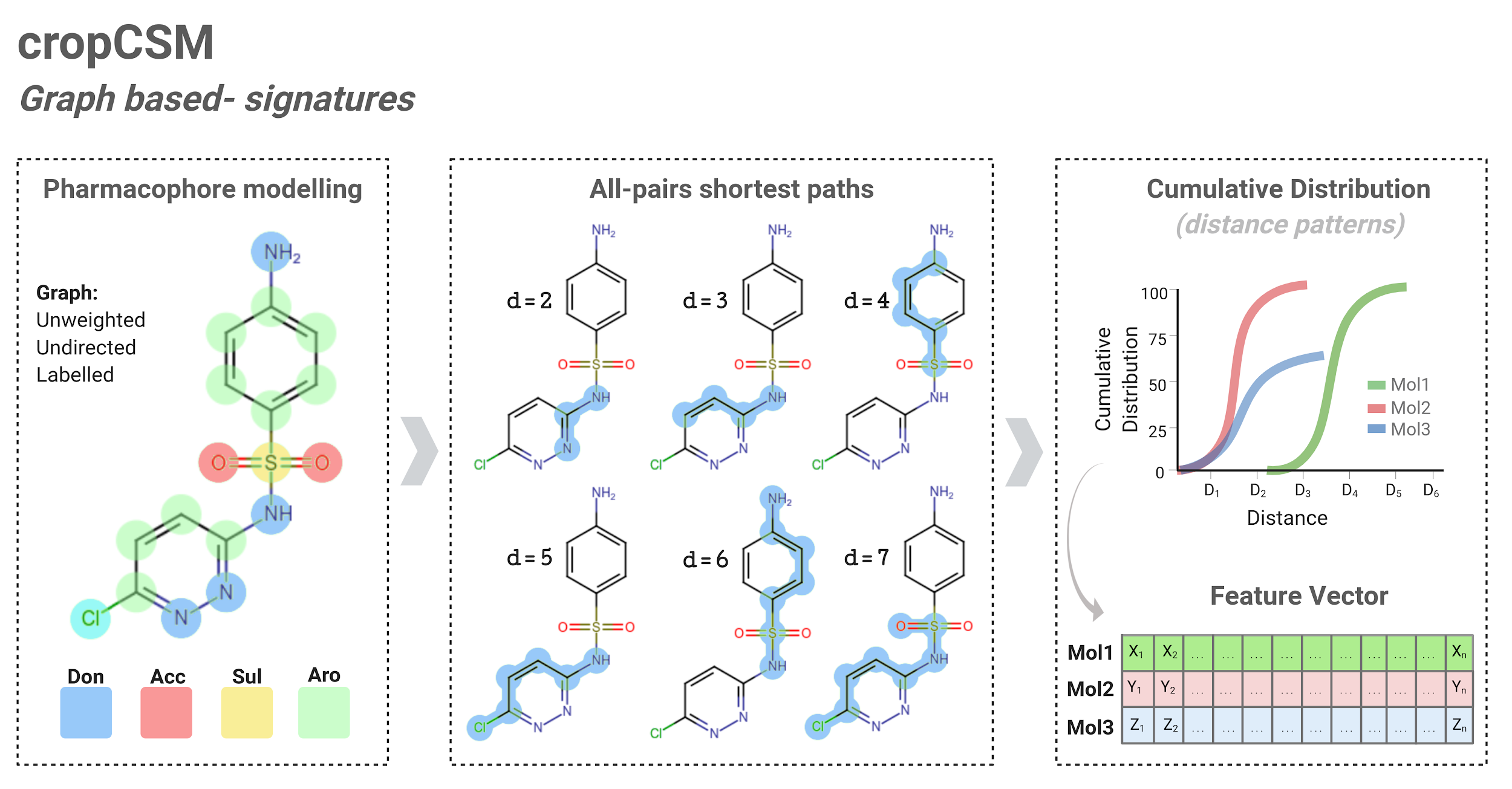

**Supplementary Fig. 3.** Modeling small molecule activity using graph-based signatures. Small molecules are modelled as unweighted, undirected graphs where nodes represent atoms and edges represent chemical bonds, with atoms labelled via pharmacophore modelling (left panel). All-pairs shortest paths are calculated between different label types (middle panel) in order to represent molecules, their geometry and physicochemical properties as cumulative distributions (right panel).

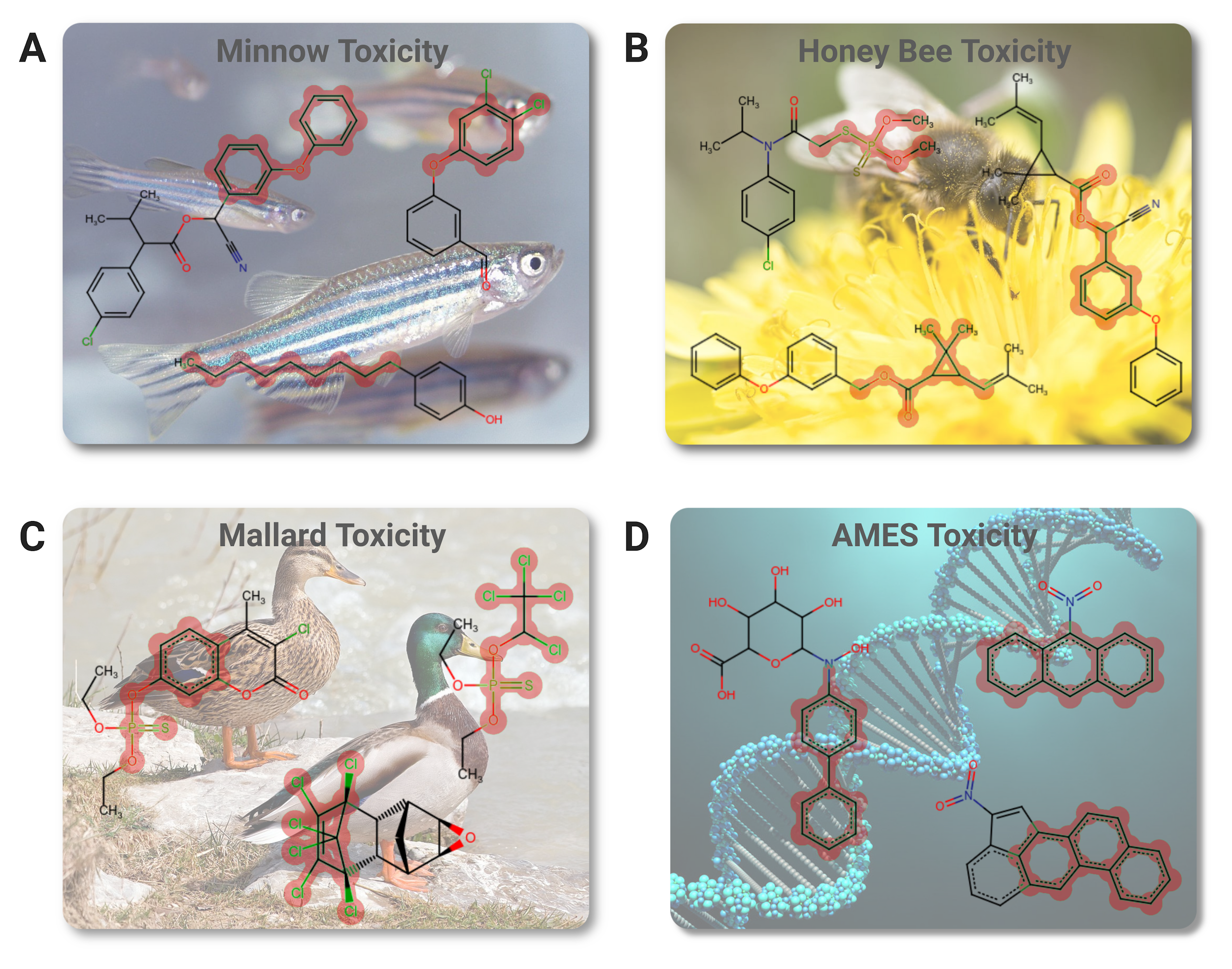

**Supplementary Fig. 4.** Substructure mining for toxicity predictors. The figure depicts common enriched substructures in compounds deemed toxic for flathead minnow (A), honey bee (B), mallard (C) and based on AMES toxicity/mutagenicity (D).

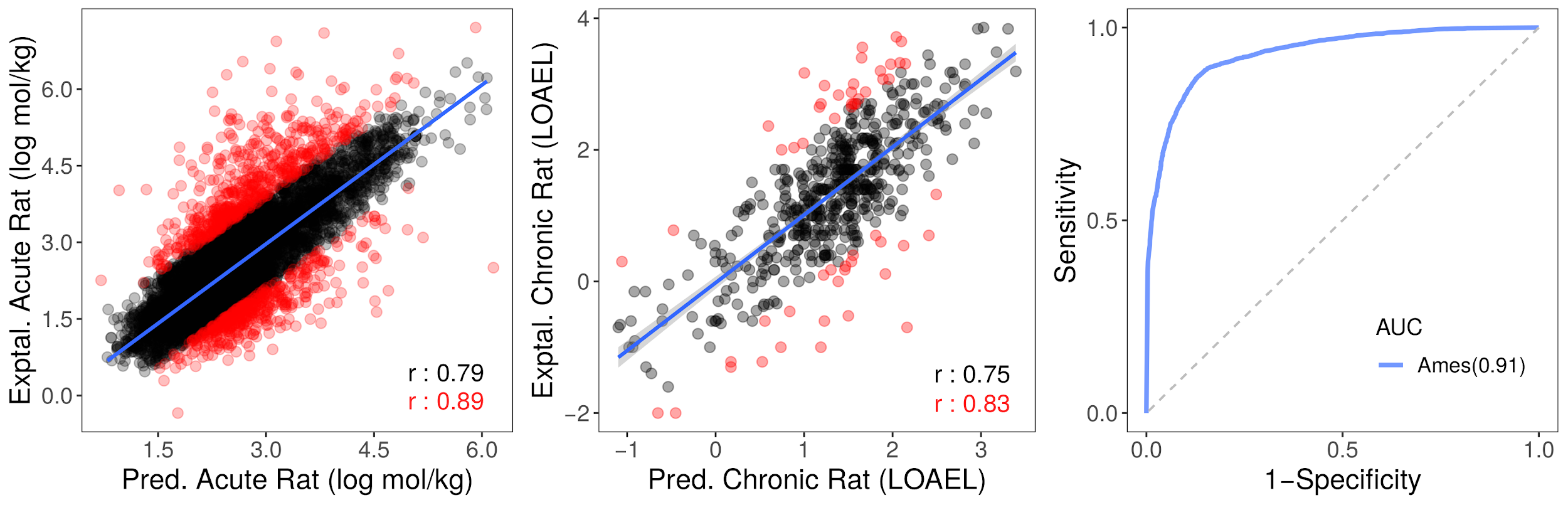

**Supplementary Fig. 5.** Performance on cross-validation of human-toxicity predictors. Three models have been developed and were capable of accurately measuring acute and chronic rat toxicity (as regression tasks, left and center graphs) as well as identifying potentially carcinogenic compounds (Ames toxicity as a classification task, right-hand side graph).

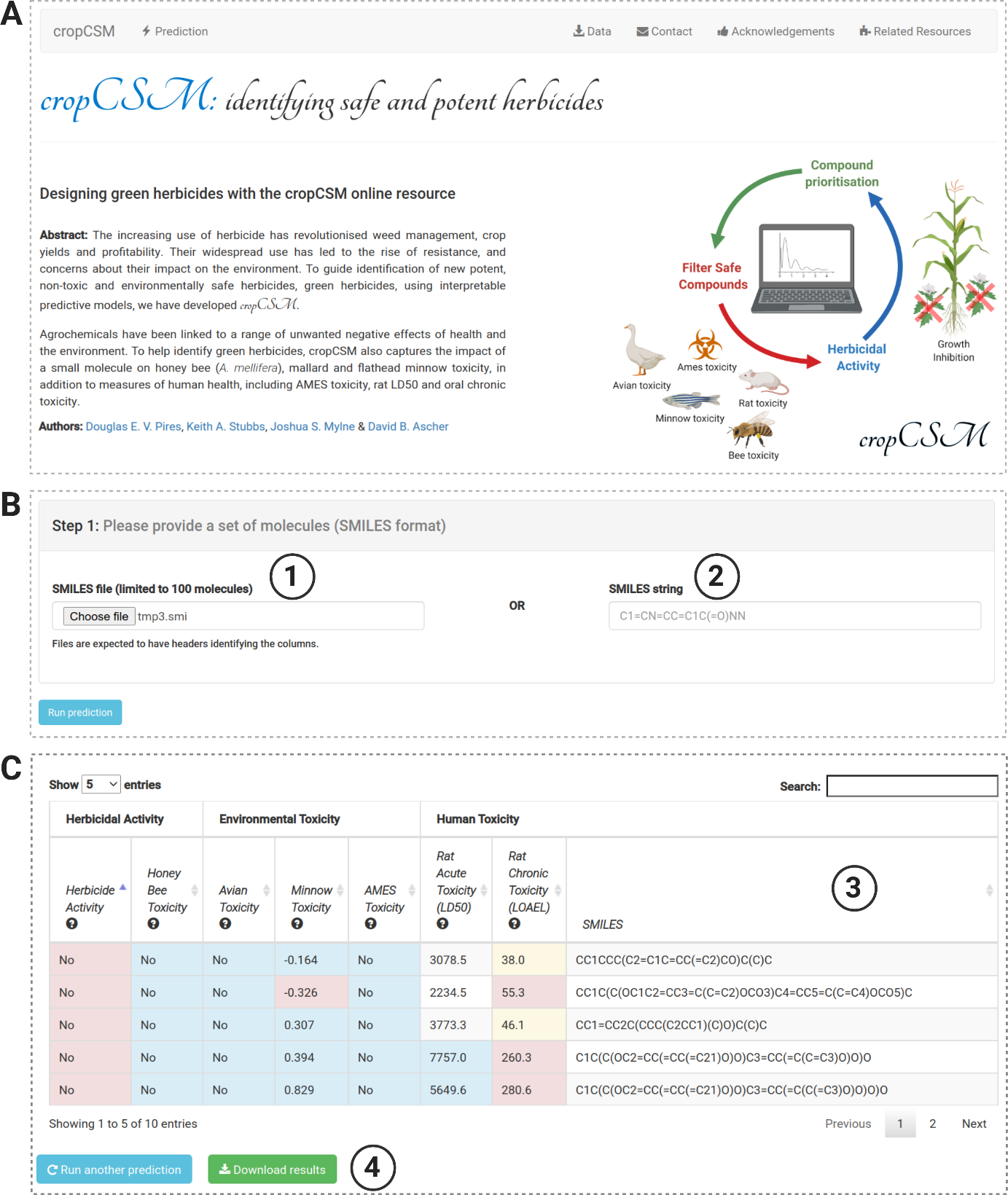

**Supplementary Fig. 6.** cropCSM web server. (A) depicts the landing page for the resource. By clicking on “Prediction” at the top menu, users are directed to the job submission page (B). There users have the options to either provide a set of molecules as a SMILES file (1) or individual molecules as a SMILES string (2). After clicking on “Run prediction”, and once calculations are complete, users are redirected to a results page (C) where predictions for herbicidal activity as well as toxicity profiles are presented in tabular format (3). Users have the options to download the results (4).
